# Widespread, depth-dependent cortical microstructure alterations in paediatric focal epilepsy

**DOI:** 10.1101/2022.11.29.518330

**Authors:** Chiara Casella, Katy Vecchiato, Daniel Cromb, Yourong Guo, Anderson M. Winkler, Emer Hughes, Louise Dillon, Elaine Green, Kathleen Colford, Alexia Egloff, Ata Siddiqui, Anthony Price, Lucilio Cordero Grande, Tobias C. Wood, Shaihan Malik, Rui Pedro A.G. Teixeira, David W. Carmichael, Jonathan O’Muircheartaigh

**Affiliations:** Centre for the Developing Brain, School of Biomedical Engineering and Imaging Sciences, King’s College London, London, UK; Department for Forensic and Neurodevelopmental Sciences, Institute of Psychiatry, Psychology and Neuroscience, King’s College London, London, UK; Department of Human Genetics, University of Texas Rio Grande Valley, Brownsville, Texas, United States; Department of Radiology, Guy’s and Saint Thomas’ Hospitals NHS Trust, London, UK; Department of Biomedical Engineering, King’s College, London, UK; Biomedical Image Technologies, ETSI Telecomunicación, Universidad Politécnica de Madrid & CIBER-BBN, ISCIII, Madrid, Spain; Department of Neuroimaging, King’s College, London, UK; MRC Centre for Neurodevelopmental Disorders, UK

## Abstract

**Objective:** Tissue abnormalities in focal epilepsy may extend beyond the presumed focus. The underlying pathophysiology of these broader changes is unclear, and it is not known whether they result from ongoing disease processes, treatment-related side-effects, or whether they emerge earlier. Few studies have focused on the period of onset for most focal epilepsies, childhood. Fewer still have utilised quantitative MRI, which may provide a more sensitive and interpretable measure of tissue microstructural change. Here, we aimed to determine common spatial modes of changes in cortical architecture in children with heterogeneous drug-resistant focal epilepsy and, secondarily, whether changes were related to disease severity.

**Methods:** To assess cortical microstructure, quantitative T1 and T2 relaxometry (qT1 and qT2) was measured in 43 children with drug-resistant focal epilepsy [age-range=4-18 years] and 46 typically-developing children [age-range=2-18 years]. We assessed depth-dependent qT1 and qT2 values across the neocortex, as well as their gradient of change across cortical depths. We also determined whether global changes seen in group analyses were driven by focal pathologies in individual patients. Finally, as a proof-of-concept, we trained a classifier using qT1 and qT2 gradient maps from patients with radiologically-defined abnormalities (MRI-positive) and healthy controls, and tested if this could classify patients without reported radiological abnormalities (MRI-negative).

**Results:** We uncovered depth-dependent qT1 and qT2 increases in widespread cortical areas in patients, likely representing microstructural alterations in myelin or gliosis. Changes did not correlate with disease severity measures, suggesting they may represent antecedent neurobiological alterations. Using a classifier trained with MRI-positive patients and controls, sensitivity was 62% at 100% specificity on held-out MRI-negative patients.

**Significance:** These findings suggest the presence of a potential imaging endophenotype of focal epilepsy, detectable irrespective of radiologically-identified abnormalities, and potentially evident pre-symptomatically.

**Key Points:**

- We assessed cortical microstructure in children with focal epilepsy
- Quantitative T1 and T2 relaxometry (qT1 and qT2) was measured in the neocortex
- Patients showed extensive qT1/qT2 increases and intracortical organization changes
- Alterations may appear during cerebral development, prior to disease onset

## 1. Introduction

Increasing evidence from neuroimaging ^1–4^, neurophysiological ^1^, neuropathological ^5^, and genetic studies ^6^, suggests that common epilepsies share disturbances in distributed cortico-subcortical brain networks ^7,8^. However, the pattern, consistency and cause of these disturbances are currently unknown, as it is uncertain how these may relate to clinical parameters such as functional decline ^9–11^.

In focal epilepsy, subtle but widespread structural abnormalities have been demonstrated in cortical areas remote from the putative epileptic focus ^12–15^. Such cortical changes may be attributed to the adverse effects of medication ^16,17^, and ongoing and cumulative disease processes ^18,19^. Alternatively, it is possible that such alterations represent a predisposing factor to the development of epilepsy. ^13,20^.

Most previous MRI studies in epilepsy have assessed structural morphological changes, including volume, surface area, and cortical thickness ^8,21^. Despite providing useful information, morphological markers can reflect the combination of several neuroanatomical features ^22^. Quantitative MRI (qMRI) techniques such as longitudinal (T1) and transverse (T2) relaxometry (qT1 and qT2) are sensitive to tissue structure and biophysical properties at the micrometre scale ^23^ and allow the in vivo assessment of inconspicuous changes in cortical tissue properties, such as myelin and free-water content ^24^.

In adults, previous research has revealed qT1 increases in ipsilateral temporal and frontal limbic cortices in temporal lobe epilepsy (TLE) patients, with more marked effects in upper cortical levels that taper off towards the white/grey matter (WM/GM) boundary. Increases were particularly marked in patients with early disease onset, suggesting that these alterations may reflect atypical neurodevelopment ^25^.

Ahmad et al. ^21^, used qT2 and surface-based analysis techniques to investigate normal-appearing cortical tissue in patients with focal cortical dysplasias (FCDs). Widespread cortical qT2 increases in frontal, parietal, and some temporal regions were observed in patients, suggesting effects of the disease in cortical regions beyond the identified focus.

There is relatively little neuroimaging research in paediatric epilepsy. Crucially, due to their shorter disease-duration, the assessment of children with epilepsy enables a deeper insight into whether cortical alterations previously reported in adults ^12,14,21,25,26^ may represent antecedent neurobiological alterations ^27^, and possibly predisposing factor to disease development ^13^, rather than a secondary effect of the disease.

A recent study ^4^ evaluated the distribution of whole-brain qT1 and qT2 changes in a sample of both children and adults with pharmaco-resistant focal epilepsy with no apparent lesions on conventional MRI. The authors reported multi-regional qT1 increases in patients, in both GM and WM ipsilateral to the epileptic origin, involving key structures of the limbic/paralimbic systems. The detected qT1 increase correlated with younger seizure-onset age, longer epilepsy duration, and higher seizure frequency, suggesting that such changes may be related to seizure burden.

In this study, we investigated surface-based qT1 and qT2 values at increasing cortical depths ^28^, as well as their gradient of change as an index of microstructural organization ^25^ in a heterogeneous sample of children with focal epilepsy. Assessing a wide range of patients with diverse underlying pathologies allowed us to get a deeper insight into whether biologically distinct focal epilepsy syndromes show robust, common microstructural deficits. As children typically have a shorter disease duration and therefore less opportunity for chronic disease-related cortical damage, we expected to detect focal, rather than global, cortical abnormalities ^25^. The relationship of these relaxation parameters to clinical factors including age of onset, disease duration and number of seizures per year was also tested.

Further, as an exploratory analysis, we evaluated the potential of qMRI features to act as a biomarker of focal epilepsy, even when there is no radiologically identified focus. For this purpose, we trained a classifier using whole-brain qT1 and qT2 cortical features from MRI-positive patients and healthy controls, and tested if the resulting classifier could identify MRI-negative patients and controls held out from the training phase.

## 2. Materials and methods

### 2.1. Subjects

Participants were recruited prospectively, and ethical approval was granted by the Health Research Authority (HRA) and Health and Care Research Wales (HCRW) (ethics ref 18/LO/1766). Infants younger than 6 months and young adults older than 18 years were excluded, as were people with major neurologic conditions unrelated to epilepsy, and contraindications for 3T MR imaging.

Eligible patients were recruited through the paediatric neurology outpatient clinics at the Evelina London Children Hospital, King’s College Hospital and Great Ormond Street Hospital, while children without epilepsy were recruited from existing volunteer databases, mainstream schools, and from the King’s College London recruiting webpage.

For participants under 16 years of age, written signed consent was obtained from the parents (or the person with legal parental responsibility) prior to any data collection. Adolescents 16 years or over provided their own written consent to participate.

A total of 96 children were recruited for the present study (ethics ref. 18/LO/1766), 89 of which were included in the analysis -43 with drug resistant focal epilepsy [age range: 4-18, mean = 12yrs, 22 female], and 46 healthy controls [age range: 2-18, mean = 11.5yrs, 27 female]. Of the 7 participants excluded from the analysis, 4 presented incidental findings and 3 had missing data. There was no significant difference in age between patients and controls (*p = 0.68*).

In patients, diagnosis and lateralization of the seizure focus were determined by a comprehensive evaluation including detailed history, neurological examination, review of medical records, video-EEG recordings, and clinical MRI evaluation. Amongst the patients assessed, the majority had temporal involvement, followed by frontal, occipital, parietal, and occipital and parietal (Table 1). A detailed description of the patient sample and their relevant clinical findings is provided in Supplementary Table 1.

**Table 1.** Descriptive demographics of the study sample.

| Characteristics | Healthy Control Participants | Patients with Epilepsy | Total |
| --- | --- | --- | --- |
| Number | 46 | 43 | 89 |
| Age at scan (years) |  |  |  |
| Mean + SD | 11.54 ± 3.96 | 12 ± 3.56 | 11.85 ± 3.76 |
| 1 - 5 | 3 | 1 | 4 |
| 6 -10 | 18 | 14 | 32 |
| 11 - 15 | 17 | 20 | 37 |
| 16 - 18 | 8 | 8 | 16 |
| Sex |  |  |  |
| Male | 19 | 21 | 40 |
| Female | 27 | 22 | 49 |
| Focus laterality (hemisphere) |  |  |  |
| Left | - | 25 |  |
| Right | - | 18 |  |
| Focus area (ratio/percentage) |  |  |  |
| Frontal | - | 13/43 |  |
| Temporal | - | 22/43 |  |
| Parietal & Occipital | - | 1/43 |  |
| Occipital | - | 3/43 |  |
| Parietal | - | 4/43 |  |
| Age of onset |  |  |  |
| Mean + SD | - | 6.95 ± 3.01 |  |
| Disease duration |  |  |  |
| Mean + SD | - | $4.85 \pm 3.16$ | |
| Number of seizures per year |  |  |  |
| Mean + SD | - | $173.46 \pm 288.08$ | |

### 2.2 Image Acquisition

Children were scanned without sedation on a 3T Achieva-TX (Philips Healthcare) using a 32-channel head coil. They were asked to stay still during scanning while watching a movie. The protocol included: a 3D T1-weighted MPRAGE [TR = 7.7ms, TE = 3.6ms, flip angle = 8°, TI = 900ms, echo-train length = 154 and acquisition time = 286s]; a T2-weighted FLAIR [TR = 5000ms, TE = 422ms, TI = 1800ms, echo-train length = 182, acquisition time = 510s]; a joint system relaxometry (JSR) acquisition ^29^, consisting of two SPGR (flip anglelJ=lJ2.2° and 12.5°, TRlJ=lJ6.2ms) and three bSSFP (with a 12° flip angle and a 180° phase increment, 49° flip angle and a 0° phase increment and 49° flip angle and a 180° phase increment respectively; all had a TR oflJ6ms). Parallel imaging acceleration (SENSE) of 1.4 was used along both phase-encoding directions. The field of view was 240 × 188 × 240mm, and the resolution was 1 mm isotropic for all images.

### 2.3. Motion Tolerance

All scans were acquired using the DISORDER scheme (Supplementary Figure 1), which has demonstrated improved tolerance against motion by guaranteeing that the acquisition of every shot contains a series of samples distributed incoherently throughout k-space ^30,31^. In-depth detail of the reconstruction algorithm has been described previously ^30,32^.

**Figure 1.**
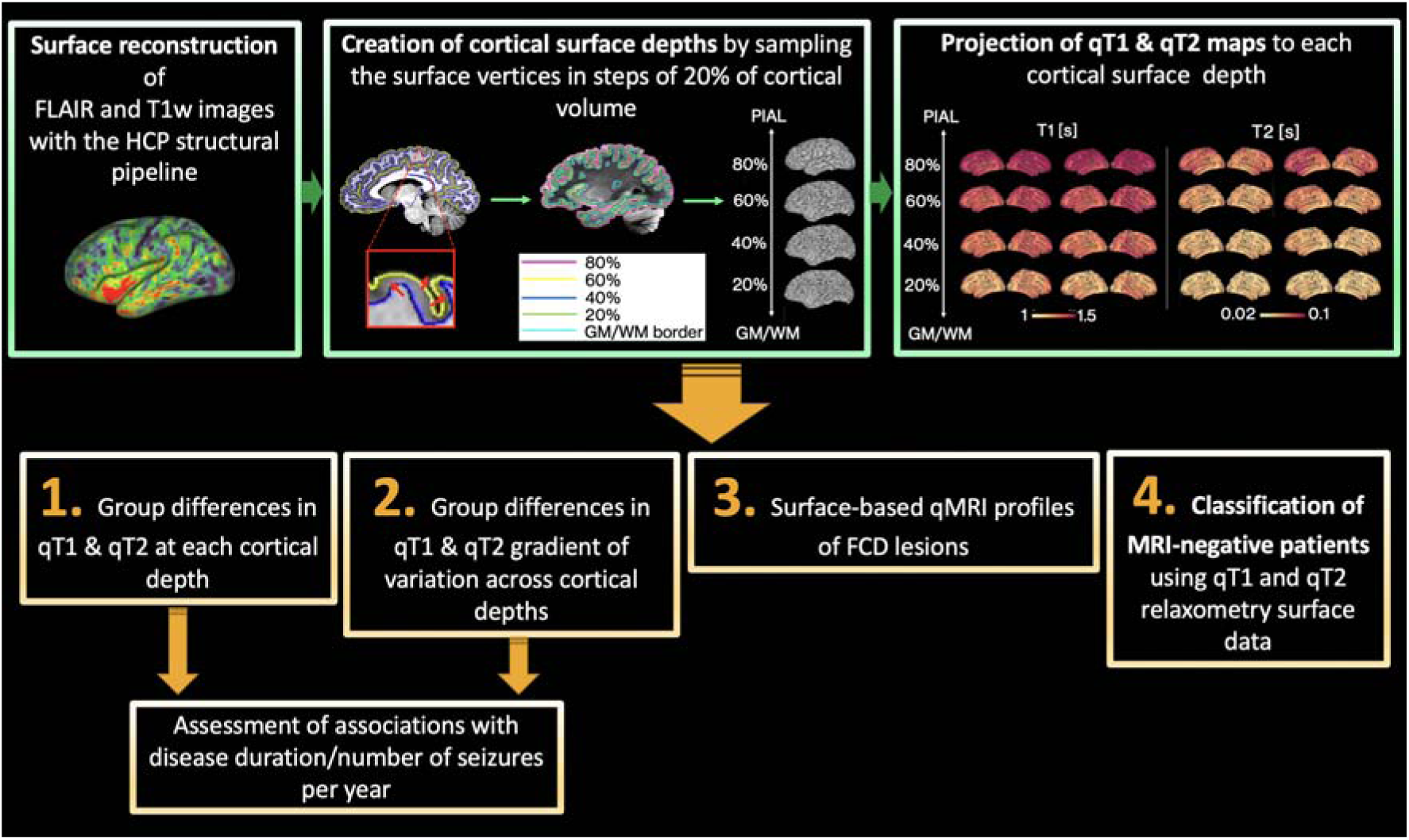
Diagram of data processing (green boxes) and analysis (orange boxes). First, surface reconstruction was performed with the HCP structural pipeline ^28^. Then, surface extraction was performed at a series of cortical depths and qT1 and qT2 maps were projected onto each depth. This allowed the assessment of group differences in: (1) qT1 and qT2 values at several depths from the pial surface; (2) qT1 and qT2 gradient of variation across cortical depths. Associations with disease duration and number of seizures per year were also explored. We also computed surface-based qMRI profiles of FCD lesions (3), and assessed whether cortical microstructure patterns learnt from MRI-positive patients could classify MRI-negative patients (4).

### 2.4. Image Analysis

#### Surface reconstruction

FLAIR and MPRAGE images were co-registered and jointly analysed with the Human Connectome Project (HCP) structural pipeline ^28^ to reconstruct WM/GM matter boundaries and pial surfaces. Full pre-processing steps and the code to run the HCP pipeline can be found at https://github.com/Washington-University/HCPpipelines.

These surfaces were used to compute equi-volume cortical surface depths by sampling surface vertices in steps of 20% of cortical volume (0%: WM/GM boundary, 100%: pial surface) across the entire cortex in Connectome Workbench (https://github.com/Washington-University/workbench).

#### T1 and T2 surface mapping

JSR ^29^ fits were performed within the QUIT toolbox ^33^, providing 3D qT1 and qT2 relaxation time maps. qT1 and qT2 images were rigidly co-registered to the FLAIR and MPRAGE volume, smoothed (s = 0.4) and sampled at each cortical depth. All volumetric images had matched distortions.

#### Between-group differences in cortical qT1 and qT2

The HCP structural pipeline outputs cortical surface metrics that are left-right symmetrical, meaning that a vertex with a specific number will refer to roughly homologous areas in each hemisphere ^28^. Therefore, we flipped the qT1 and qT2 surface maps of patients with right hemispheric focus so that they could be analysed with patients with left hemispheric focus. This enabled us to both maximize sample size and to better understand the impact of focus laterality on detected changes.

Group-wise differences at 20%, 40%, 60% and 80% cortical distance from the GM/WM border to the pial cortical surface were tested. Additionally, at each vertex, qT1 and qT2 values at 20% distance were subtracted from those at 80% distance, providing a gradient of relaxation values across the cortex, here an index of intracortical organisation. Group differences in this gradient were tested. Finally, associations between qT1 and qT2 changes in patients with age of disease onset, disease duration and number of seizures per year were tested. Because of skewness in the data, the number of seizures per year data were log-transformed before assessing correlations. All models were tested by permutation (using the permutation analysis of linear models package, PALM ^34^).

To understand whether the focal cortical *location* had a significant impact on any observed changes, the above analyses were then repeated by splitting patients with temporal focus (N = 22) and patients with frontal focus (N = 13) into two sub-groups. Here, we excluded patients with occipital (N =3) and parietal (N = 4) foci, because of low numbers, as well as those patients who presented with both occipital and temporal foci (N = 1).

As a first step, for all analyses we tested for an interaction between age and group. Where this was not significant, age as well as sex were thus regarded as covariates of no interest in the final analyses. Vertex-wise cortical thickness was also controlled for in the analyses, because of evidence that this feature is altered in epilepsy ^12,15^. Finally, differences in cortical curvature have been shown to systematically affect the laminar and myeloarchitecture of the cortex ^36^, so that strong correlations between local curvature and R1 (1/T1) can be observed. Therefore, we also controlled for individual differences in vertex-wise cortical curvature. To address the multiple comparisons problem, family-wise error correction rate was applied across MRI modalities (i.e. qT1 and qT2) and statistical contrasts using the “-corrmod” and “-corrcon options” in PALM. Threshold-free cluster enhancement (TFCE) was employed as test statistic ^35^, and relationships were tested by permutation using the Draper-Stoneman method ^34^, which ensures that spatial dependencies in the data are preserved for all permutations.

#### Surface-based qMRI profiles of individual focal cortical dysplasias (FCDs)

To assess whether changes seen in group differences may have been driven by focal lesions in individual patients, we performed a post-hoc examination of the qMRI profiles of lesions of 5 patients with radiologically-suspected FCDs. For this purpose, we first manually delineated lesion masks on the FLAIR images in each patient. We then mapped the lesion masks to each cortical depth in both patients and controls in the HCP standard surface space, consisting of approximately 32,000 vertices (Figure 2) ^28^. Finally, we extracted the unpermuted residuals of qT1 and qT2 values at different cortical depths within the lesion area. These were computed as follows: Y-Z*g, where Z is the matrix with confounds (i.e. age, sex, thickness and curvature) and g are the estimated GLM regression coefficients from Y = X*b + Z*g.

**Figure 2.**
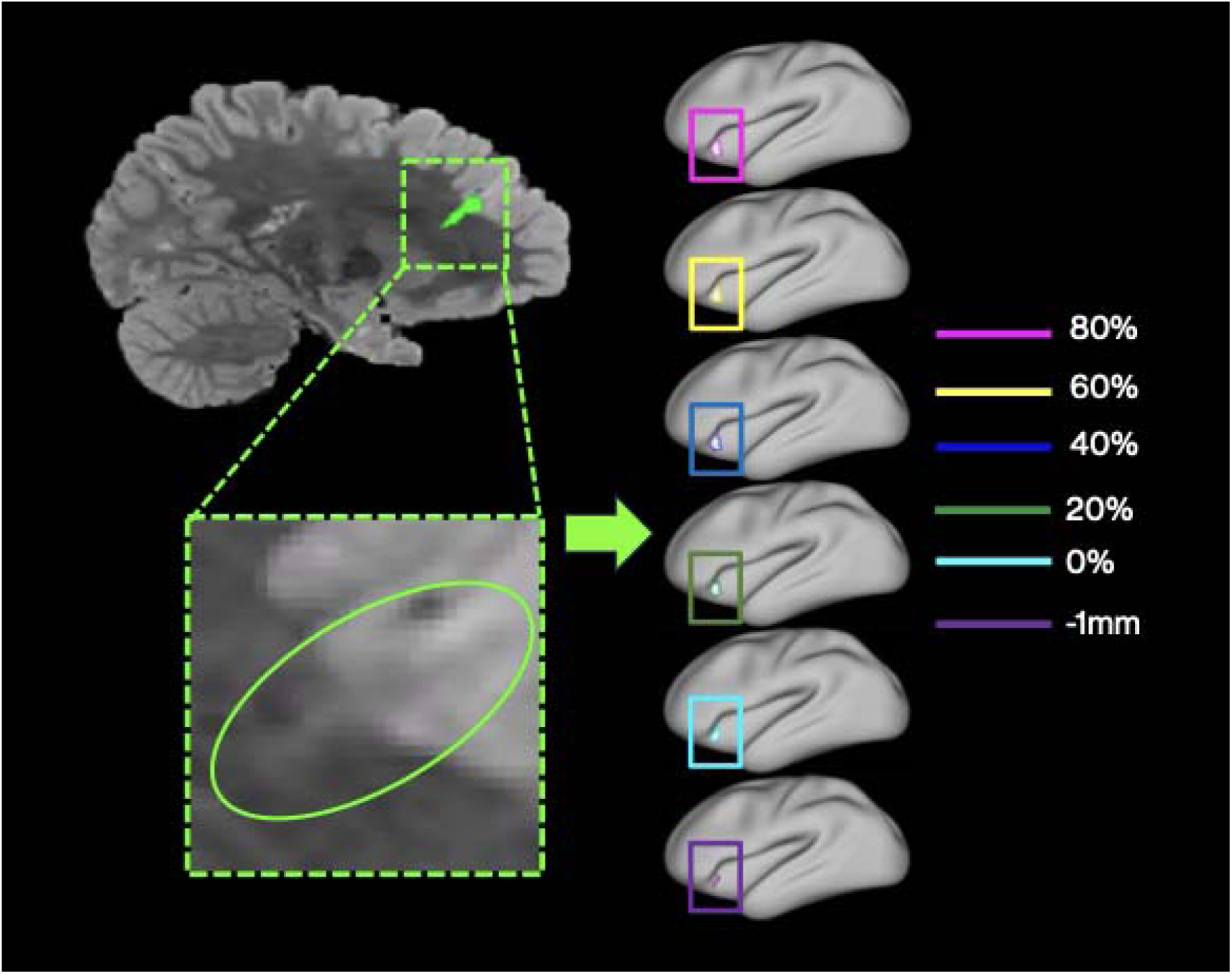
Workflow of the computation of surface-based qMRI profiles of FCD lesions. Lesion masks were delineated on the FLAIR images of each of the 5 patients with FCDs. The lesion masks were then mapped to each cortical depth in patients and controls in standard surface space. The mean and standard deviation of qT1 and qT2 were extracted within the masks, after removing the effect of age and sex using linear regression. This allowed computation of the z-score profile of each patient normed against the controls, corrected for age and sex.

This allowed computation of the z-score profile of each patient normed against the controls, corrected for age and sex. Importantly, for this part of the analysis, we further extended the spatial sampling to 1mm below the GM/WM border to increase sensitivity to transmantle signs.

#### Testing diagnostic performance of surface qMRI features

Using Scikit-learn ^37^, a random forest classifier ^38^ was trained on the gradient maps of qT1 and qT2 MRI-positive patients (N = 17) and controls (N = 31), after removing the effect of age and sex using linear regression. Both qT1 and qT2 features for the whole brain were concatenated and input to the classifier. After training, the model’s performance was tested on qT1 and qT2 gradient maps MRI-negative patients (N = 26) and controls (N = 14) held-out from the training set.

For completeness, the same procedure was then applied to train a classifier on the data of MRI-negative patients (N = 26) and controls (N = 14). This was then tested on MRI-positive patients (N = 17) and controls (N = 31) held-out from the training set.

For both models, area under the ROC Curve (AUC), accuracy, sensitivity, specificity and F1lJscores were used to evaluate classification performance. A permutation-based p-value was calculated to assess the probability that the observed results could have been obtained by chance.

## 3. Results

### 3.1. Group differences in qT1 an qT2 at each depth from the grey/white matter border

#### Cohort Differences

Increases in qT2 at the 60% and 80% distance from the GM/WM border were detected in patients. Although the effects were stronger in the left hemisphere, increases in qT2 were spread across the two hemispheres. Additionally, increases in qT1 were detected at 60 and 80% depth from the GM/WM border in the ipsilateral hemisphere in patients (Figure 3A, top). Figure 3C (left quadrant) plots subject-average qT1 and qT2 values at 80% distance for patients and controls in areas where significant group differences were detected, against age.

**Figure 3.**
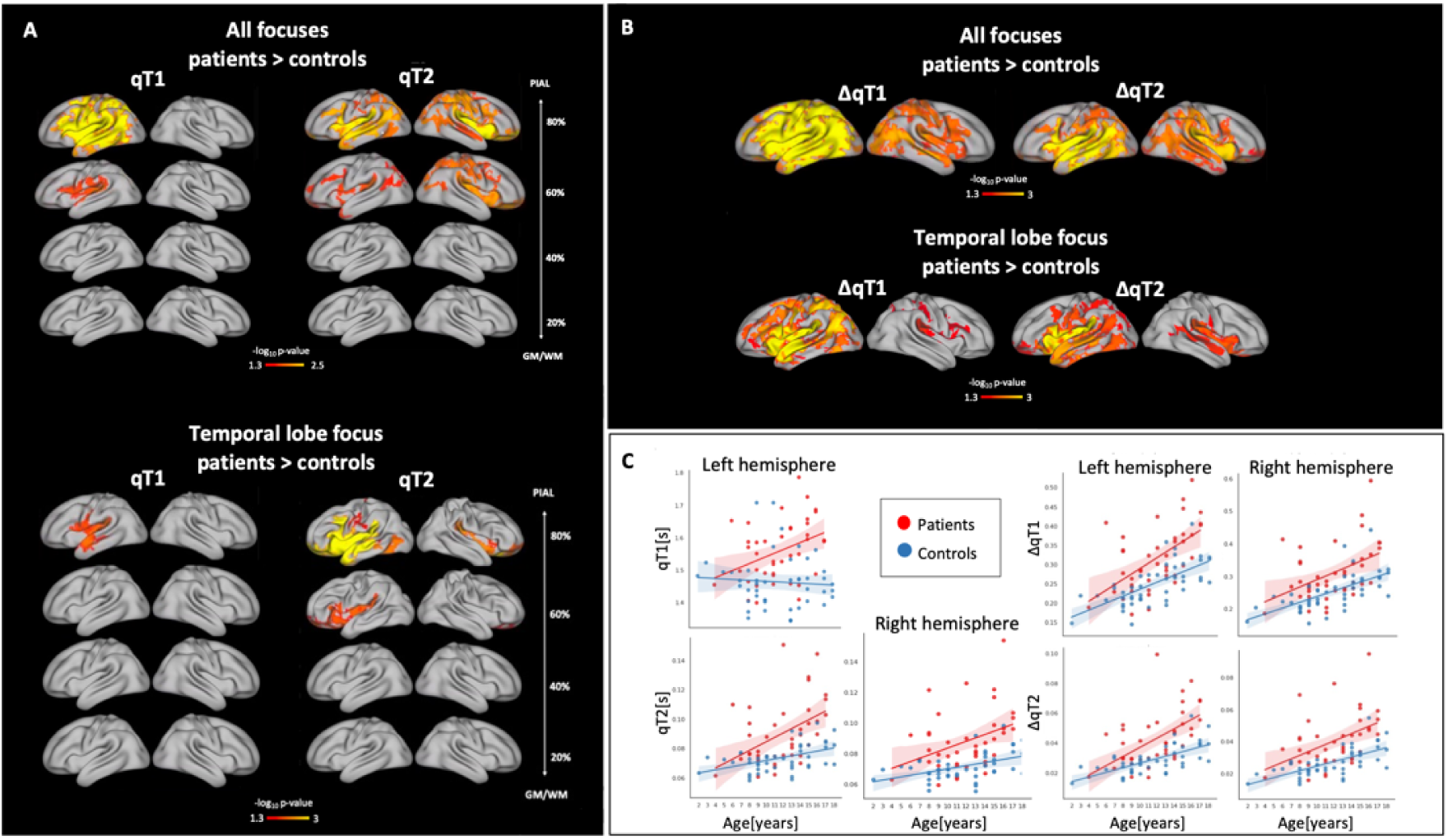
(A) Group differences between patients and healthy controls in qT1 and qT2 at each cortical depth: When comparing all patients against controls, qT1 increases were detected in the ipsilateral hemisphere at 60 and 80% depth in patients, while bilateral increases were detected in qT2 at 60 and 80% depth. When splitting patients into frontal versus temporal lobe focus, qT1 increases were detected in the left temporal lobe in TLE patients compared to controls. Additionally, qT2 increases were detected at 60 and 80% distance in TLE patients compared to controls. **(B) Group differences in qT1 and qT2 gradients (**Δ**qT1 and** Δ**qT2):** When comparing all patients against controls, widespread steeper qT1 and qT2 variation was detected in patients across both hemispheres. When splitting patients into subgroups based on focus, widespread steeper qT1 and qT2 variation was detected in TLE patients compared to controls. Alterations extended beyond the temporal lobe. **(C) Relationship between age and average qT1 and qT2 values in patients and controls**: On the left quadrant, average qT1 and qT2 values at 80% distance from the GM/WM border are plotted for each subject against age in areas where significant group differences were detected. On the right, average qT1 and qT2 gradient values are plotted against age for each subject.

#### Lobar Differences

When splitting patients into frontal and temporal lobe groups, qT1 increases were detected at 80% distance from the GM/WM border in the left temporal lobe of TLE patients versus controls. Additionally, qT2 increases were detected at 60 and 80% distance from the GM/WM border in TLE patients, especially in the left hemisphere. Though stronger effects were detected in the temporal lobe, increases in qT2 were widespread, especially at 80% distance from the GM/WM border. We detected no significant differences between frontal patients and controls, nor between the two patients’ sub-groups (Figure 3A, bottom).

With both analytic approaches, we did not detect any association between microstructure changes and any clinical variables. However, there was a positive relationship between disease duration and age in patients (*rho = 0.59, p < 0.001*), potentially indicating the presence of collinearity, which in turn might have impacted our sensitivity to associations between microstructural changes and clinical variables.

### 3.2. Group differences in qT1 an qT2 gradients as an index of intracortical organization

A “surface-out” gradient, driven from the WM/GM border, of increasing qT1 and qT2 values could be seen across the cortex in both patients and controls.

#### Cohort Differences

Permutation tests revealed a significant difference between the two groups in cortical qT1 and qT2 gradients, indicating a steeper gradient of variation across cortical depths in the patient group, with increasingly higher qT1 and qT2 values in the outermost cortical depths across the two hemispheres (Figure 3A, bottom). The effects were widespread and bilateral, however they were stronger in the ipsilateral hemisphere. Figure 3C (right quadrant) plots subject-average qT1 and qT2 gradient values for patients and controls in areas where significant group differences were detected, against age.

#### Lobar Differences

After splitting patients based on temporal versus frontal focus, we detected steeper variation of qT1 and qT2 values across cortical depths in patients with TLE. The changes concerned mostly the left hemisphere, which was the hemisphere affected in most TLE patients. They extended beyond the temporal lobe, affecting most of the cortex. We detected no significant differences between frontal patients and controls, nor between the two patients’ sub-groups, and no significant associations with clinical variables.

### 3.3. Surface-based qMRI profiles of FCD lesions

Figure 4 illustrates the surface maps of the focal cortical dysplasia (FCD) locations, as well as the z-score profiles of FCD lesions across cortical depths, for each patient. The detected deviations from the control distribution were heterogeneous between lesions and mostly not significant, suggesting that spatial modes of qT1 and qT2 alterations are not driven by focal abnormalities.

**Figure 4.**
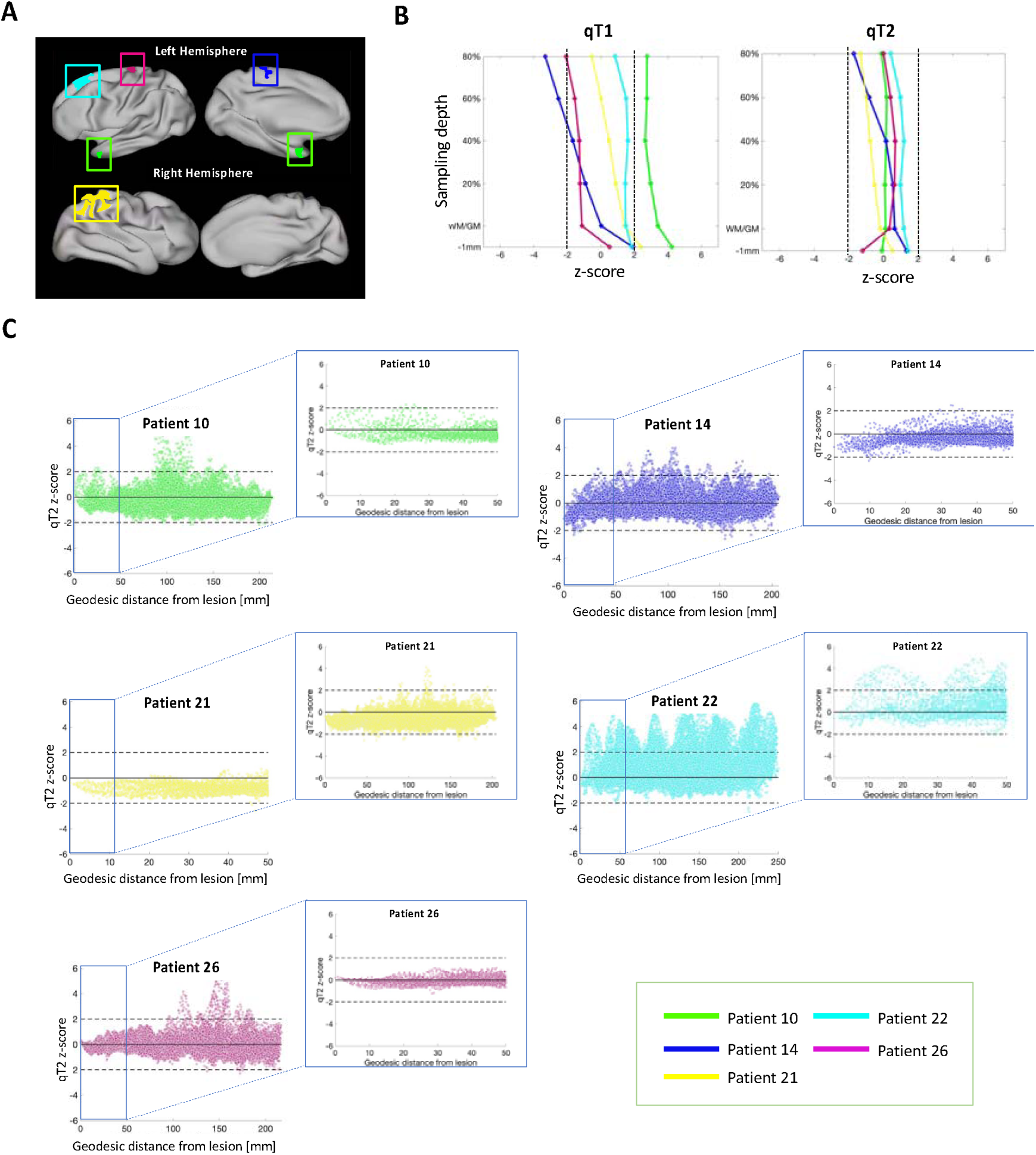
Spatial modes of qT1 and qT2 alterations are not driven by focal abnormalities. (A) Surface maps of FCDs locations of all patients. Lateral and medial views are presented for both hemispheres. The FCDs were located in temporal, frontal and parietal areas. (B) qMRI z-score profiles of FCDs for qT1 (left) and qT2 (right). Here, each line represents the z-score deviation of one patient against all controls, across cortical depths. Any z-score greater than +2 or smaller than -2 (black dashed lines) represents a significant deviation from the control distribution. Detected changes were heterogeneous between lesions and mostly not significant. (C) qT2 z-score values as a function of geodesic distance from the lesions’ centre at 80% cortical depth. Both a cortex-wise and a zoomed-in view (i.e. within 50mm of the lesion centre) are plotted. In all patients the spread of abnormalities presents an “FCD-fugal” pattern, with abnormalities affecting cortical areas remote from the FCD.

### 3.4. Classifying epilepsy in MRI-negative patients with qMRI features

On a classifier trained with MRI-positive patients and controls, we had a test AUC of 90.7%, accuracy of 80%, sensitivity of 71.4% at 89.4% specificity, and an F1 score of 79% on MRI-negative patients and controls that were not part of the training group. To test robustness to training data, we flipped the training and test datasets. When the classifier was trained with MRI-negative patients and controls, we had a test AUC of 90.4%, accuracy of 74%, sensitivity of 60% at 87.5% specificity, and an F1 score of 69% on MRI-positive patients and controls that were not part of the training group. The classification performance could not be attributed to chance [*p = 0.001*] (Figure 5).

**Figure 5.**
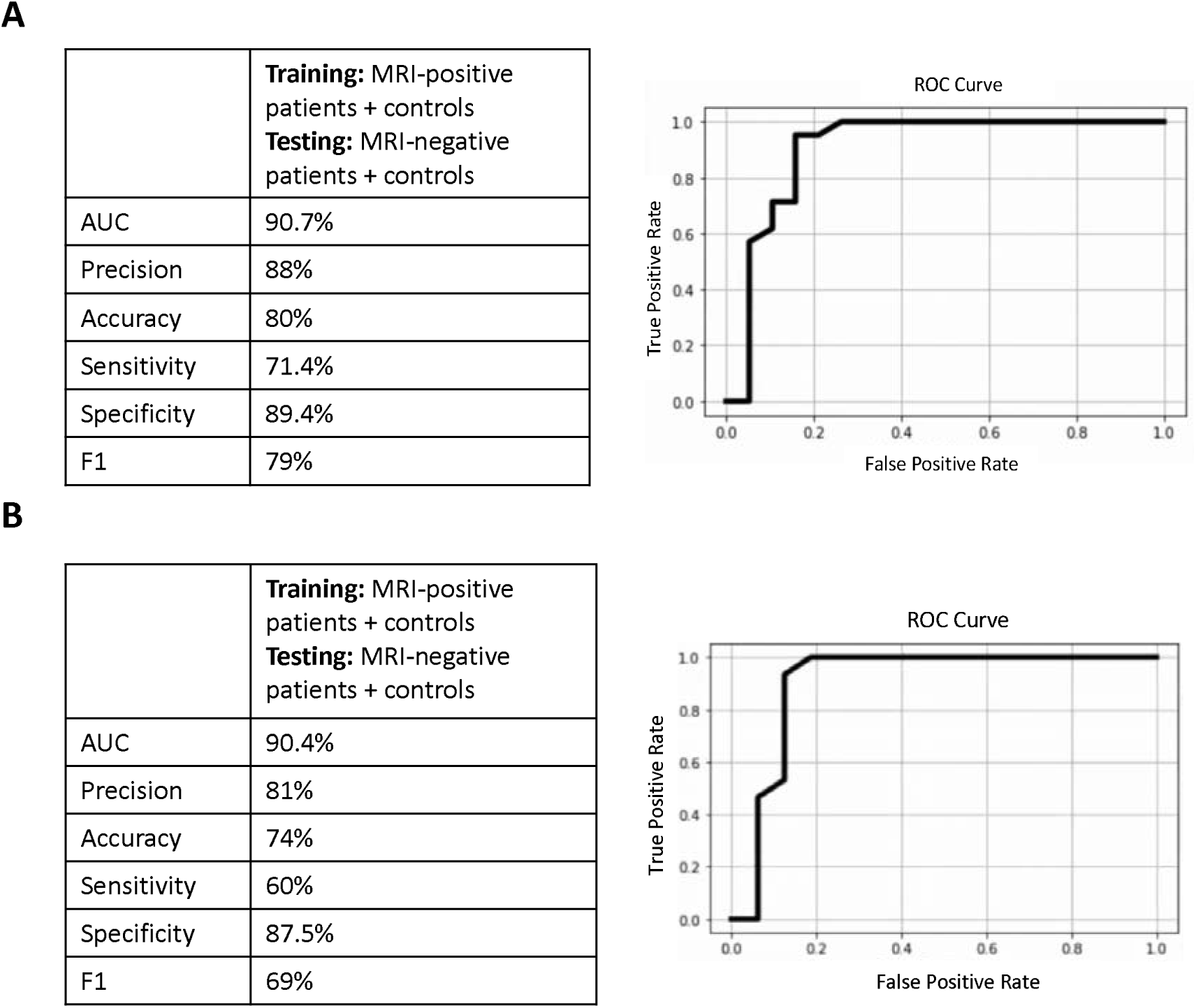
Random forest classifiers performance. For both classifiers, the confusion matrix and ROC curve are displayed. For both classifiers the potential for classification could not be attributed to chance and the p-value was significant.

## 4. Discussion

Increasing evidence suggests that biologically distinct epilepsy syndromes may be characterized by shared disturbances in cortico-subcortical brain networks ^7,8^ and, specifically in focal epilepsies, cortical abnormalities extend spatially beyond the presumed epileptogenic focus ^4^. In this work, we assessed whole brain cortical changes in children with drug-resistant focal epilepsy.

Focussing exclusively on childhood enabled us to gain a deeper insight into whether cortical alterations are the result of ongoing chronic disease-processes ^4,18,19^ and treatment-related side-effects ^16,17^, or whether they emerge at an earlier disease-stage ^13,27^. The marked heterogeneity in the sample in terms of lesion/focus location and underlying pathologies could be leveraged to further understand shared patterns of abnormalities. The use of a multiparametric qMRI approach allowed to move beyond morphological markers to probe changes in cortical tissue microstructure ^24^.

Overall, our findings indicate the presence of consistent, widespread, depth-mediated qT1 and qT2 increases in children with focal epilepsy. Additionally, they reveal significant alterations in patients’ intracortical organization ^39^, demonstrated by a radial gradient of increased qT1 and qT2 values driven from the WM/GM border in patients. Consistent with previous evidence ^21,25,40^, the observed cortical changes were robust to potential confounders, including age-at-scan and brain morphology, suggesting that they represent a different underlying pathophysiological mechanism. Moreover, the detected alterations did not appear specific to, or driven by, the focal cortical lesions of patients, as demonstrated by the fact that qT1 and qT2 values within the investigated FCDs did not significantly deviate from control values in the same area. Importantly, in accord with previous studies ^12,21^, clinical parameters did not correlate with qT1 and qT2 times across the cortex, implying that such microstructural alterations may be independent of ongoing disease processes.

Notably, our findings indicate that microstructural alterations in focal epilepsy are independent of seizure onset or focus laterality. Though stronger changes were detected in the temporal lobe of TLE patients, alterations still extended to the wider cortex. Such widespread effects were particularly evident when looking at qT1 and qT2 gradients. While no alterations were detected in the sub-group of patients with frontal focus, this is most likely because of the smaller size of this subgroup. Additionally, while qT1 increases at the different depths predominantly affected patients’ ipsilateral hemisphere, we detected bilateral effects on the qT1 gradient over cortical depths. Finally, extensive qT2 increases were uncovered in patients across cortical depths and when looking at the gradient of change. Our findings are in accord with previous work demonstrating the presence of morphological alterations following a bilateral pattern and extent that is independent of the side of seizure focus ^3^, and not specific to the patients’ focal pathology ^3,4^.

Importantly, we demonstrate that these widespread cortical features are sufficiently consistent across all children with epilepsy that they can act as classification features. A simple random forest classifier trained on whole-brain qT1 and qT2 surface maps from MRI-positive patients and controls could classify epilepsy status in MRI-*negative* patients and controls not used in the training phase. Likewise, the same classifier trained on whole-brain qT1 and qT2 surface maps from MRI-negative patients and controls could classify epilepsy status in MRI-*positive* patients and controls not used in the training phase. These findings suggest that there may be an imaging endophenotype of focal epilepsy, common in MRI-positive and MRI-negative cases.

There are several underlying microstructural changes that could explain the cortical qT1 and qT2 alterations observed in patients in this study. For qT1, previous evidence suggests that this measure may be sensitive to GM myelin content ^41^. Therefore, increased qT1 values and alterations in qT1 gradients may potentially reflect the presence of cortical myelin disturbance in paediatric focal epilepsy, mirroring pathology findings in TLE where surgical specimens had alterations in intracortical myelination and fiber arrangement, particularly in upper cortical layers ^42^.

On the other hand, the qT2 depth-wise and gradient increases observed in this study might be related to gliosis. Previous evidence has reported a correlation between gliosis and hippocampal subfields qT2 in TLE ^43^, though this is not yet reported in cortical epilepsy. Therefore, gliosis on a microstructural level may reflect a remodelling process which might explain the observed qT2 increases ^21^. Alternatively, since qT2 is dependent on the free water content in tissue ^24,44^, tissue damage and replacement of cells in nervous tissue by water could potentially be another mechanism leading to the increased cortical qT2 observed in patients.

From an aetiological point of view, ongoing seizure-activity could underly qT1 and qT2 alterations observed in this study. Accordingly, Su and colleagues (2023) reported limbic/paralimbic qT1 increases in children and adults with focal epilepsy which correlated with younger seizure-onset age, longer epilepsy duration, and higher seizure frequency, suggesting again that such changes may be related to seizure burden ^4^.

However, other previous investigations ^12,21^ failed to detect a significant relationship between structural parameters and clinical seizure occurrence, and reported the lack of an association between anticonvulsive drugs and qT2 variation ^21^. Crucially, the ability of our classifier to classify young patients, as well as the lack of association between qT1 and qT2 changes and disease duration or age of onset demonstrated in this study, lend support to the suggestion that alterations may be independent of disease progression and instead might constitute antecedent microstructural alterations reflecting developmental cortical changes ^20,25^.

Further supporting the hypothesis that qT1 and qT2 abnormalities reflect precursor anomalies in brain development, a longitudinal study of children with new-onset focal and idiopathic generalized epilepsy demonstrated differences in baseline GM and WM volumes of focal patients compared to controls ^46^. Further, Widjaja and colleagues reported changes in cortical thickness in the right medial temporal, cingulate, and left frontal cortices in children with new-onset seizures ^27^.

It is possible that such early brain changes may predispose patients to seizures, but this remains a challenging hypothesis to test in focal epilepsy compared to family designs possible in genetic epilepsies ^47,48^. Future relaxometry studies assessing paediatric patients with new-onset seizures longitudinally will enable to better disentangle the impact of age from the effect of disease, as well as of recurrent seizures and long-term antiepileptic medications on the brain. Additionally, the specificity of these global cortical changes to focal epilepsy as opposed to seizures is unclear and inclusion of other epilepsies is needed.

Future studies could also assess subtle variations in cortical iron depositions and myelination, thus providing rich complementary information to the existing evidence. Additionally, the integration of multimodal MRI with macroscale diffusion-based connectivity changes promises to better understand both the local-and network-level substrates of drug-resistant focal epilepsies.

Most importantly, analysis approaches that address patients heterogeneity, such as normative modelling ^50^, will allow to control for brain variability and rapid, non-linear tissue changes associated with development, and therefore to better disentangle significant pathologic changes from ‘normal’ or expected age-related alterations.

## 5. Conclusion

To conclude, we demonstrate widespread, depth-mediated qT1 and qT2 increases in the cortex of children with treatment-resistant focal epilepsy. Though the aetiology of these cortical alterations is not fully understood, it is possible that such microstructural changes appear during cerebral development and represent precedent neurobiological alterations, rather than the cumulative effect of recurrent seizures or the consequence of medications’ side-effects. Accordingly, based on our findings it is plausible to suggest that global cortical changes in surface-based qT1 and qT2 may constitute a potential imaging endophenotype of focal epilepsy, detectable with or without the presence of radiologically identified abnormalities and disease-onset.

## CRediT Statement

**Chiara Casella:** Conceptualisation, Data Curation, Formal Analysis, Investigation, Methodology, Project Administration, Software, Visualisation, Writing - Original Draft Preparation; Writing - Review & Editing; **Katy Vecchiato:** Conceptualisation, Data Curation, Formal Analysis, Investigation, Methodology, Project Administration, Writing - Original Draft Preparation; **Daniel Cromb:** Project Administration; **Yourong Guo:** Software; **Anderson M. Winkler:** Software, Validation; **Emer Hughes:** Investigation, Project Administration; **Louise Dillon:** Investigation, Project Administration; **Elaine Green:** Investigation, Project Administration; **Kathleen Colford:** Investigation, Project Administration; **Alexia Egloff:** Investigation, Project Administration; **Ata Siddiqui:** Investigation, Project Administration; **Anthony Price:** Project Administration, Resources, Software; **Lucilio Cordero Grande:** Resources, Software; **Tobias C. Wood:** Resources, Software; **Shaihan Malik:** Resources; **Rui Pedro A.G. Teixeira:** Resources, Software; **David W. Carmichael:** Conceptualisation, Formal Analysis, Methodology, Resources, Supervision, Writing - Review & Editing; **Jonathan O’Muircheartaigh:** Conceptualisation, Formal Analysis, Funding Acquisition, Methodology, Resources, Supervision, Writing - Review & Editing.

## Conflict of Interest/Ethical Publication Statement

We confirm that we have read the Journal’s position on issues involved in ethical publication and affirm that this report is consistent with those guidelines.

## Acknowledgements

We are grateful to the families who generously supported this study. We also thank the Pediatric Neurology team from the Evelina London Children Hospital, including Dr. Ruth Williams, Dr. Elaine Hughes, Dr. Shan Tang, and Dr. Karine Lascelles.

