## Supplementary figures and images for "Widespread, depth-dependent cortical microstructure alterations in paediatric focal epilepsy"

### Supplementary Figure 1

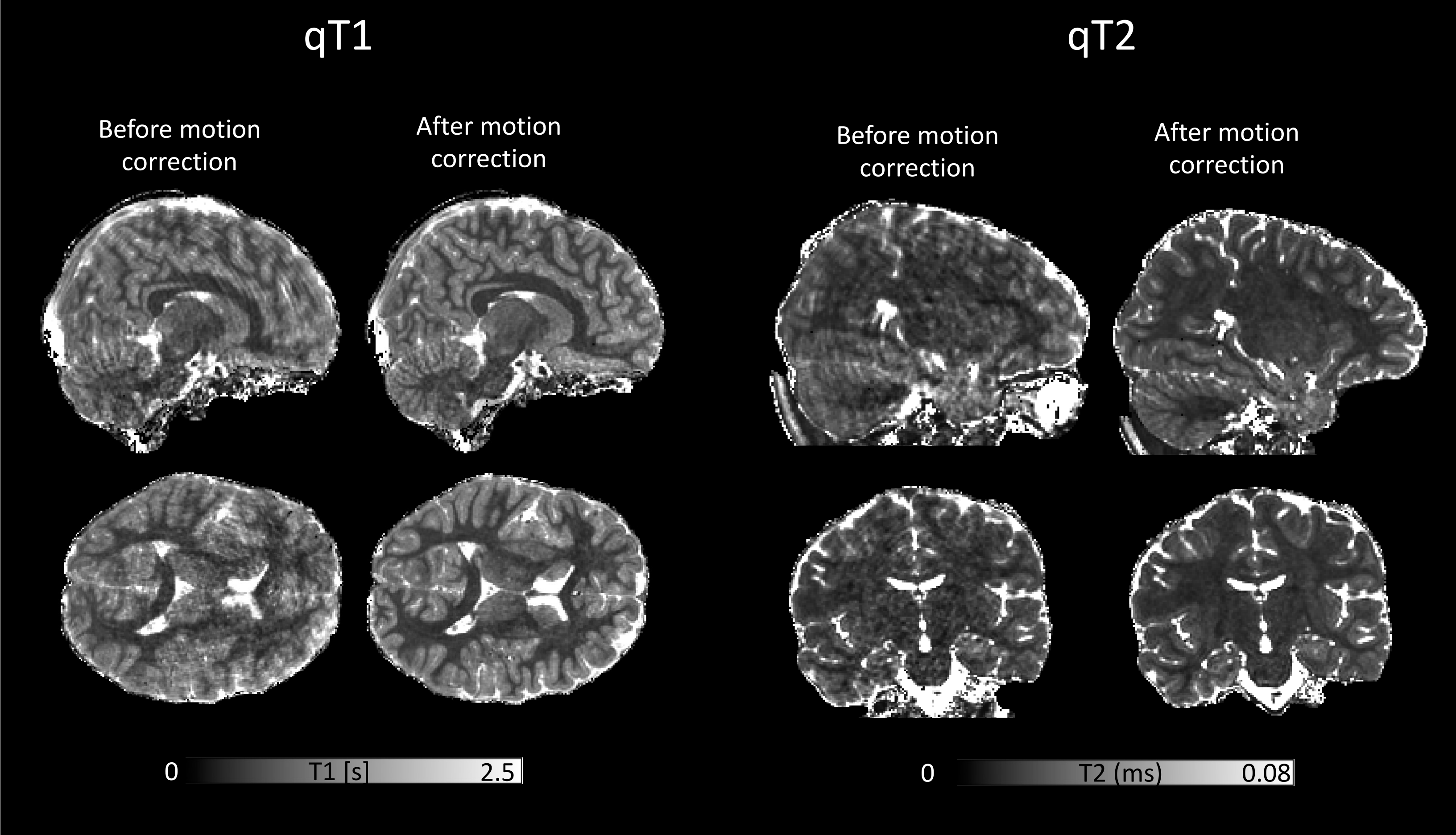
