## Supplementary Table 1 for "Widespread, depth-dependent cortical microstructure alterations in paediatric focal epilepsy"

***Supplementary Table 1. Clinical details of the patients assessed in this study.***

| **Patient** | **Age of onset** | **Length of disease** | **Number of seizure in the last year** | **Secondary generalized tonic-clonic seizures** | **Current medication** | **EEG** | **PET** | **3T MRI** | **MRI report summary** | **Comorbidities** |
| --- | --- | --- | --- | --- | --- | --- | --- | --- | --- | --- |
| 1 | 7 | 2 | 30 | No | Carbamazepine | Right fronto-temporal | N/A | Right temporal | Right hippocampus: bulkier and hyperintense T2 with subtle loss of internal architecture. |  |
| 2 | 4 | 7 | 100 | No | Carbamazepine, Lamotrigine, Melatonin | Left frontal | Left frontal | Negative | No clear localizing or lateralizing features appreciated. |  |
| 3 | 7 | 1 | 100 | No | Carbamazepine, Levetiracetam, Phenobarbital | Left frontal | Left temporal and secondary left inferior frontal | Negative | Mild prominence of the ventricle and sulcal spaces without any interval atrophy. |  |
| 4 | 5 | 9 | 100 | Yes | Lamotrigine, Lacosamide, Topiramate | Left parietal | Bilateral | Negative | Aspecific white matter signal change bilaterally at the tips of the temporal horns, unrelated to epilepsy. |  |
| 5 | 11 | 1 | 100 | No | Levetiracetam | Left occipital | Left temporal | Negative | Ventricular asymmetry, left occipital horn more prominent than right. | Developmental delay, speech delay, dyspraxia |
| 6 | 3 | 3 | Unknown | Yes | Levetiracetam, Sodium valproate | Left frontal | Left frontal | Negative | Minor asymmetry in the parietal lobes but with no obvious cortical changes. |  |
| 7 | 10 | 4 | 104 | Yes | Topiramate, Lacosamide | Right frontal | Right frontal | Right frontal | Epilepsy surgery with incomplete resection prior to MRI scan: some high FLAIR signal at the margins of the resection cavity but findings are unspecific. |  |
| 8 | 6 | 6 | 28 | No | Lamotrigine, Clobazam | Right temporal | N/A | Negative | No clear localizing or lateralizing features appreciated. |  |
| 9 | 4 | 6 | 0 | No | Ethosuximide, Lacosamide | Left frontal | N/A | Negative | No clear localizing or lateralizing features appreciated. |  |
| 10 | 5 | 7 | 100 | No | Topiramate, Perampanel, Sertraline | Left temporal | Right temporal | Left temporal | **Suspected FCD**  Slightly increased signal intensity in the left temporal lobe, with some blurring of the GM/WM interface. | Asperger syndrome, speech delay, ADHD, developmental delay |
| 11 | 5 | 4 | 200 | No | Carbamazepine, Lamotrigine, Clozapam | Left frontal | Normal | Negative | No clear localizing or lateralizing features appreciated. |  |
| 12 | 6 | 6 | 999+ | No | Carbamazepine, Topiramate, Clobazam | Left temporal | Left temporal | Negative | No clear localizing or lateralizing features appreciated. | Hypothyroidism |
| 13 | 9 | 3 | 0 | No | Levetiracetam Lamotrigine | Left frontal | N/A | Right frontal | No clear localizing or lateralizing features appreciated. | Neonatal injury |
| 14 | 3 | 6 | 100 | No | Carbamazepine, Lacosamide, Clobazam | Left parietal | N/A | Left parietal | **Suspected FCD**  Cortical thickening and abnormal gray-white matter differentiation in left mesial frontoparietal region. This is associated with abnormal cortical and subcortical white matter signal with tapering towards the ventricle. |  |
| 15 | 5 | 5 | 999+ | No | Carbamazepine, Lamotrigine, Sodium Valproate | Right frontal | Bilateral parietal | Negative | No clear localizing or lateralizing features appreciated. |  |
| 16 | 11 | 5 | 100 | No | Carbamazepine, Perampanel | Multiple EEGs are normal and fail to capture any episodes | N/A | Negative | No clear localizing or lateralizing features appreciated. |  |
| 17 | 7 | 5 | 200 | No | Sodium Valproate | Left occipital | Left parietal, left occipital | Negative | No clear localizing or lateralizing features appreciated. |  |
| 18 | 3 | 12 | 200 | No | Sodium Valproate, Topiramate | Left temporal | Bilateral temporal | Negative | No clear localizing or lateralizing features appreciated. | Autism spectrum disorder, speech delay, developmental delay, dyslexia |
| 19 | 7 | 1 | 342 | Yes | Carbamazepine, Levetiracetam Sodium Valproate | Right parietal, right occipital | N/A | Right occipital | Grey matter heterotopia. |  |
| 20 | 1 | 11 | 60 | No | Carbamazepine, Lacosamide | Left temporal, left parietal | N/A | Negative | No clear localizing or lateralizing features appreciated. |  |
| 21 | 8 | 2 | 999+ | No | Oxcarbazepine | Right parietal | Right frontal | Right parietal | **Suspected FCD**  Asymmetry in the parietal lobes with a deeper right intraparietal sulcus/ Large area of cortical-subcortical mildly increased T2 signal with blurring of the GM/WM junction involving predominantly the parietal lobe. | Developmental delay, anxiety, depression |
| 22 | 12 | 3 | 39 | Yes | Levetiracetam, Sodium valproate | Left frontal, left temporal | N/A | Left frontal | **Suspected FCD**  Left frontal lobe area of abnormal cortico-subcortical signal. |  |
| 23 | 5 | 3 | 0 | No | Levetiracetam | Right temporal | N/A | Right temporal | Gyral crowding in the right inferolateteral temporal lobe. |  |
| 24 | 9 | 3 | 4 | No | Carbamazepine, Lamotrigine | Right temporal | N/A | Negative | No clear localizing or lateralizing features appreciated. | Autism spectrum disorder, anxiety |
| 25 | 9 | 7 | 0 | No | Carbamazepine | Right parietal, right occipital | N/A | Right occipital | Asymmetric signal intensity in the subcortical white matter adjacent to the tip of the right occipital jorn and calcarine sulcus with subtle blurring of the GM/WM interface. | Developmental delay |
| 26 | 4 | 4 | 100 | No | Carbamazepine, Levetiracetam | Left parietal | Left parietal | Left parietal | **Suspected FCD**  Subtle left medial precentral gyrus cortical high signal. |  |
| 27 | 5 | 12 | 2 | Yes | Lamotrigine, Sodium valproate | Right temporal | N/A | Right temporal, right parietal | Encephalomalacia of right caudate head and right occipital gliosis. | Right neonatal stroke, speech delay, developmental delay, learning disabilities, depression, anxiety disorder, ADHD, disrupted sleep |
| 28 | 10 | 5 | 20 | Yes | Carbamazepine, Sodium Valproate | Right temporal | Right temporal | Negative | No clear localizing or lateralizing features appreciated. |  |
| 29 | 4 | 6 | 50 | Yes | Carbamazepine | Left frontal, left parietal | N/A | Negative | No clear localizing or lateralizing features appreciated. |  |
| 30 | 10 | 7 | 14 | Yes | Carbamazepine, Levetiracetam | Left temporal | N/A | Left temporal | Asymmetry of the amygdala/hippocampus. |  |
| 31 | 9 | 5 | 14 | No | Levetiracetam, Lacosamide | Bilateral frontal and temporal | Left frontal, left temporal | Negative | No clear localizing or lateralizing features appreciated. |  |
| 32 | 2 | 4 | 170 | No | Lamotrigine, Sodium Valproate | Left temporal | N/A | Negative | No clear localizing or lateralizing features appreciated. |  |
| 33 | 4 | 0.5 | 999 | No | Carbamazepine | Right frontal | Right temporal/Right parietal | Right frontal | Abnormal GM/WM differentiation noted on FLAIR but not clearly seen in other sequences – could be artefactual. |  |
| 34 | 7 | 3 | 50 | No | Oxcarbazepine | Right temporal | N/A | Negative | No clear localizing or lateralizing features appreciated. |  |
| 35 | 9 | 1 | 300 | Yes | Oxcarbazepine, Lacosamide | Left temporal | Left temporal | Left temporal | Abnormal subcortical high T2 in the left temporal lobe, some gyral/cortical thinning, slight sulcal widening over the left parietal convexity.  Appearances suggest an underlying genetic/autoimmune cause. | Obsessive compulsive disorder |
| 36 | 4 | 13 | 300 | Yes | Levetiracetam, Sodium valproate, Lacosamide | Left central | Left fronto-central | Negative | No clear localizing or lateralizing features appreciated. |  |
| 37 | 5 | 2 | 20 | Yes | Levetiracetam, Lamotrigine | Left temporal | N/A | Negative | No clear localizing or lateralizing features appreciated. |  |
| 38 | 7 | 3 | 6 | No | Cabarmazepine | Right Frontal | N/A | Negative | No clear localizing or lateralizing features appreciated. |  |
| 39 | 5 | 10 | 25 | No | Levetiracetam, Brivaracetam | Right frontal | Right frontal and temporal | Negative | No clear localizing or lateralizing features appreciated. |  |
| 40 | 10 | 5 | 60 | Yes | Oxcarbazepine, Lamotrigine | Left temporal | Left temporal | Negative | Smaller Left Hippocampus but no sclerosis. |  |
| 41 | 15 | 1 | 9 | Yes | Levetiracetam | Left temporal | N/A | Right parietal | Ganglioneuroma grade 1 confirmed following right parietal craniotomy. |  |
| 42 | 6 | 8 | 999+ | No | Carbamazepine, Zonisamide | Right frontal | Left temporal | Negative | No clear localizing or lateralizing features appreciated. |  |
| 43 | 1 | 14 | 120 | No | Lacosamide | Left temporal | Left temporal | Left temporal | Abnormal signal loss of GM/WM differentiation in the left mesial/anterior temporal lobe.  Hypothesised FCD or low grade tumour. | Speech delay |
